# Systematic characterization of cancer-associated SPOP mutants reveals novel and reprogrammable degradative activities

**DOI:** 10.1101/2025.11.21.689851

**Authors:** Alana G. Caldwell, Harshil Parmar, Xiaokang Jin, Chen Zhou, Xiaoyu Zhang

**Affiliations:** Interdisciplinary Biological Sciences Graduate Program, Northwestern University, Evanston, Illinois 60208, United States; Department of Chemistry, Northwestern University, Evanston, Illinois 60208, United States; Chemistry of Life Processes Institute, Northwestern University, Evanston, Illinois 60208, United States; Robert H. Lurie Comprehensive Cancer Center, Northwestern University, Chicago, Illinois 60611, United States; Center for Human Immunobiology, Northwestern University, Chicago, Illinois 60611, United States; International Institute for Nanotechnology, Northwestern University, Evanston, Illinois 60208, United States

**Keywords:** E3 ligase, Recurrent mutation, Proteomics, Protein-protein interactions, Targeted protein degradation

## Abstract

Speckle-type POZ protein (SPOP) functions as the substrate adaptor of the Cullin3-RING ligase (CRL3) complex and is recurrently mutated in multiple cancer types. Among these, F102C and F133L are frequent prostate cancer mutations within the substrate-binding domain, yet their biochemical consequences remain incompletely understood. Using quantitative proteomics, we show that SPOP-F133L, unlike SPOP-F102C, retains degradative activity toward the nuclear basket proteins NUP153 and TPR, indicating substrate-dependent loss-of-function. Moreover, SPOP-F133L induces partial down-regulation of p53 through a CRL-dependent, post-translational mechanism, revealing a potential neo-substrate relationship. Finally, we demonstrate that both SPOP-F102C and SPOP-F133L support targeted protein degradation (TPD) in an engineered cellular system. These findings define the degradative capacities of SPOP mutants and highlight opportunities to repurpose these variants as mutant-selective E3 ligases for therapeutic applications.

## Introduction

The ubiquitin-proteasome system (UPS) is the primary pathway for the degradation of intracellular proteins, thereby maintaining proteome homeostasis^[1]^. Substrate specificity within this pathway is conferred by E3 ubiquitin ligases, which recruit target proteins to ubiquitination machinery^[2]^. Among these, the Cullin-RING ligases (CRLs) represent the largest E3 family. They are modular assemblies comprised of 1) a Cullin scaffold, 2) a RING-box protein such as Rbx1 that recruits E2 ubiquitin-conjugating enzymes, and 3) a substrate-recognition receptor^[3]^. Speckle-type POZ protein (SPOP) serves as the substrate-recognition component of the Cullin3-RING ligase (CRL3) complex. The BTB domain of SPOP anchors it to the Cullin scaffold, whereas the MATH domain recognizes substrates, promoting their ubiquitination and degradation by the proteasome^[4]^ (**Figure S1A**). SPOP localizes to nuclear speckles through higher-order oligomerization, forming subnuclear hubs that promote efficient ubiquitination^[5]^. Functionally, SPOP is recognized as a tumor suppressor, as many of its established substrates are oncogenic regulators, such as BET family proteins (e.g., BRD2, BRD3, BRD4)^[6]^, the androgen receptor (AR)^[7]^, and the androgen signaling co-activator TRIM24^[8]^.

Recurrent loss-of-function mutations in SPOP have been identified in multiple cancers, most prominently in prostate cancer, where such mutations occur in up to ∼15% of primary and metastatic cases^[9]^. These mutations predominantly cluster within the substrate-binding MATH domain, frequently occurring at residues Y87, F102, M117, W131, and F133^[10]^. Among them, F102C and F133L are among the most frequently observed SPOP mutations (**Figure 1A**). These mutations disrupt substrate binding, resulting in stabilization of oncogenic targets and promotion of tumorigenesis^[10-11]^. Conversely, context-dependent oncogenic functions of SPOP have been observed in other cancer types. For example, the SPOP-M35L mutant identified in hepatocellular carcinoma increases affinity for the tumor suppressor IRF2BP2, leading to its enhanced ubiquitination and degradation, thereby promoting cell proliferation and metastasis^[12]^. Despite these insights, a comprehensive and proteome-wide understanding of how recurrent SPOP mutations alter substrate engagement remains incomplete. In this study, we employ quantitative proteomics to systematically characterize the endogenous substrates and neo-substrates perturbed by SPOP-F102C and SPOP-F133L. Moreover, we investigate whether recurrent SPOP mutations can be exploited as mutant-selective E3 ligases for targeted protein degradation (TPD) applications.

**Figure 1.**
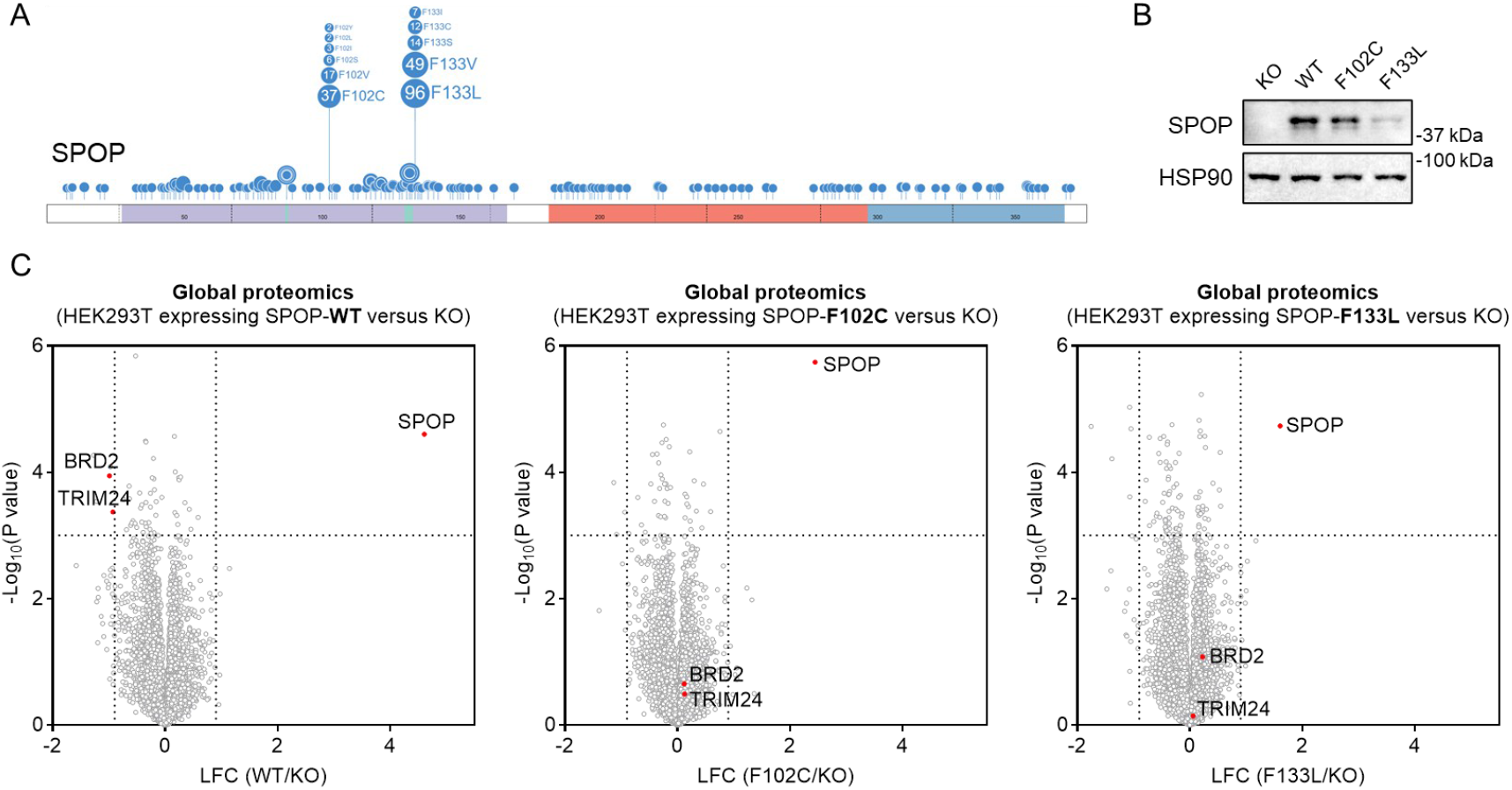
Generation of *SPOP* KO HEK293T cell models reconstituted with SPOP-WT, F102C, or F133L. **A**. Domain architecture of SPOP illustrating recurrent cancer-associated mutations clustered within the substrate-binding MATH domain. The F102C and F133L variants examined in this study are indicated. The schematic was generated using ProteinPaint (https://proteinpaint.stjude.org/). **B**. Western blot analysis of SPOP expression in *SPOP* KO HEK293T cells and in KO cells stably expressing SPOP-WT, F102C, or F133L.The result is representative of two experiments (n = 2 biologically independent samples). **C**. Volcano plot showing global proteomic changes between *SPOP* KO HEK293T cells and KO cells stably expressing SPOP-WT, F102C, or F133L (n = 4 biological independent samples). *P* values were calculated by two-sided t-test and adjusted using Benjamini-Hochberg correction for multiple comparisons.

## Results and Discussion

### Quantitative proteomic profiling of protein expression changes driven by SPOP-WT, F102C, and F133L

Previous proteomic studies have primarily focused on identifying endogenous SPOP substrates and determining whether cancer-associated SPOP mutants exhibit loss-of-function in degrading these substrates^[13]^. Given that SPOP mutations may both impair degradation of native substrates and promote degradation of neo-substrates, we sought to systematically characterize these effects using mass spectrometry-based whole-proteome analysis. We first generated *SPOP* knockout (KO) HEK293T cells (**Figure S1B**). Consistent with the established role of SPOP in targeting BRD2 and TRIM24 for degradation^[6, 8]^, both proteins were up-regulated in *SPOP* KO cells, confirming functional disruption of SPOP in this cell model (**Figure S1C** and **Table S1**). We then reintroduced SPOP wild-type (WT) and two recurrent prostate cancer mutants, F102C and F133L, into the KO background to generate stable cell lines (**Figures 2A, B**). Proteomic profiling revealed that only SPOP-WT, but not the F102C or F133L mutants, restored degradation of BRD2 and TRIM24 (**Figure 2C** and **Table S2**). These findings are consistent with previous studies indicating that both mutations are loss-of-function variants^[13c, 14]^. Thus, our reconstituted system provides a functional platform to further investigate substrate specificity and potential neo-substrate recognition by SPOP mutants.

**Figure 2.**
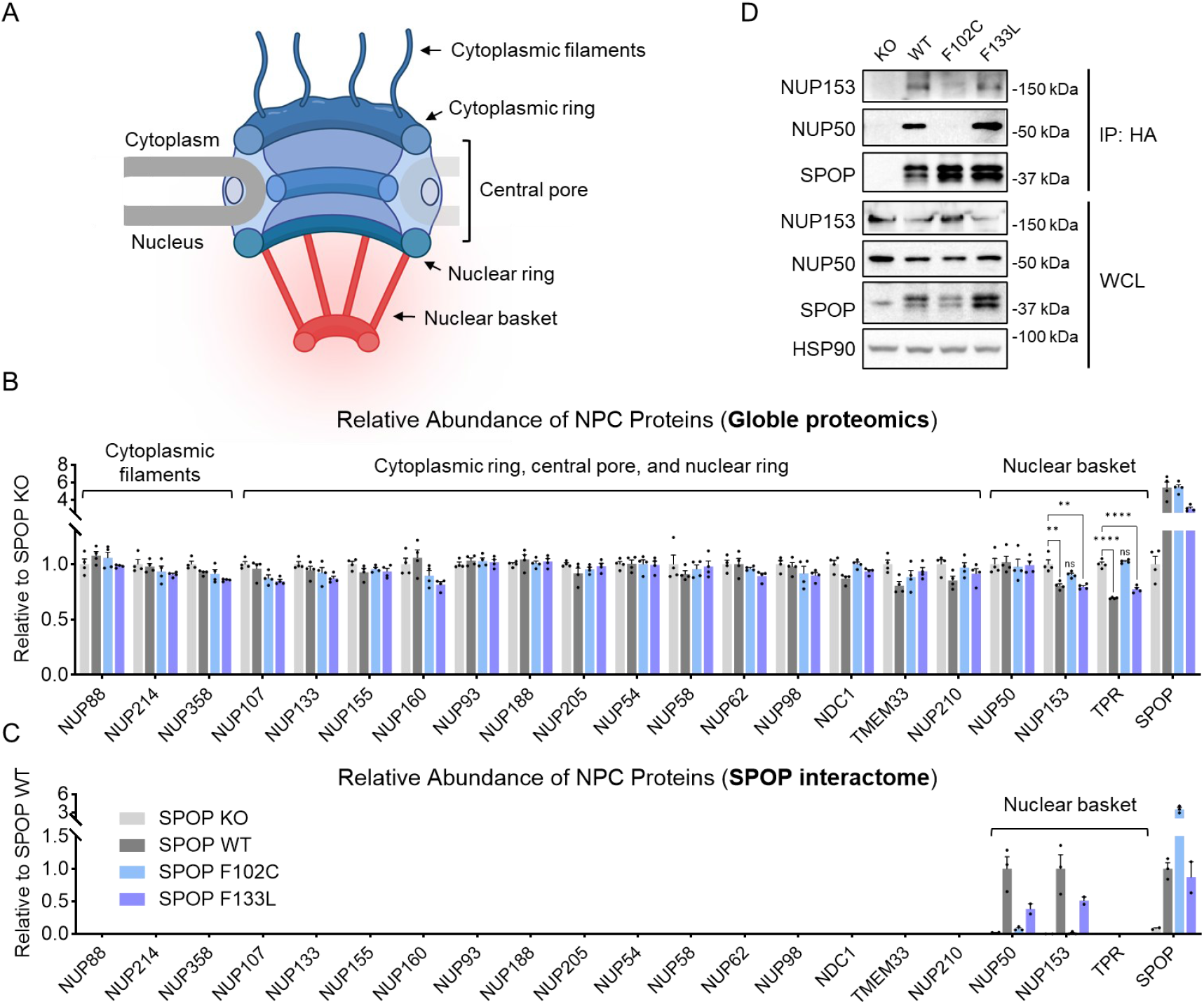
SPOP-F133L, but not SPOP-F102C, retains degradative activity toward nuclear basket proteins NUP153 and TPR. **A**. Schematic illustration of the nuclear pore complex showing the cytoplasmic, central, and nuclear basket regions. The schematic was created with BioRender.com. **B**. Quantitative proteomic analysis of 20 nucleoporins in *SPOP* KO HEK293T cells reconstituted with SPOP-WT, F102C, or F133L. Data are presented as mean ± SEM (n = 4 biological independent samples). Statistical significance was assessed using unpaired two-tailed Student’s t-tests. The color legend for each bar is shown in Figure 2C. **C**. AP-MS analysis of SPOP-WT, F102C, and F133L. Data are presented as mean ± SEM (n = 2 biologically independent samples for SPOP KO and F133L, n = 3 biologically independent samples for SPOP-WT and F102C). **D**. Co-immunoprecipitation analysis showing the interaction between HA-tagged SPOP and NUP153 or NUP50. The result is representative of two independent experiments (n = 2 biologically independent samples).

### SPOP-F133L retains degradative activity toward nuclear basket proteins NUP153 and TPR

The human nuclear pore complex (NPC) is a large macromolecular assembly composed of a number of distinct proteins known as nucleoporins^[15]^ (**Figure 2A**). A previous study identified NUP153 as a substrate of SPOP, suggesting a role for SPOP in regulating NPC function and potentially protein transport through the nuclear envelope^[16]^. The study further showed that the SPOP-F102C mutant loses the ability to degrade NUP153. Using our established cell models, we sought to investigate whether SPOP targets additional components of the NPC and whether SPOP-F133L similarly acts as a loss-of-function mutant unable to degrade nucleoporins.

Our whole-proteome analysis quantified 20 nucleoporins, containing all major NPC substructures, including the cytoplasmic filaments, cytoplasmic ring, central pore, nuclear ring, and nuclear basket (**Figure 2B** and **Table S2**). Among these, the nuclear basket protein NUP153, which is a known SPOP substrate, was reduced upon SPOP-WT expression (**Figure 2B**). Notably, the abundance of another nuclear basket component, TPR, was also reduced under the same conditions (**Figure 2B**), indicating that TPR is a potential new substrate of SPOP. In fact, NUP153 and TPR were the only nucleoporins among all quantified proteins that were down-regulated in cells expressing SPOP-WT. This suggests that SPOP may specifically regulate components of the nuclear basket within the NPC. Consistent with previous findings, the SPOP-F102C mutant lost the ability to degrade these nuclear basket proteins (**Figure 2B** and **Table S2**). Surprisingly, SPOP-F133L retained degradative activity toward both NUP153 and TPR (**Figure 2B** and **Table S2**), indicating that it may be a substrate-dependent loss-of-function mutant. These results also suggest that in cancer cells harboring the SPOP-F133L mutation, the expression levels of NUP153 and TPR may remain largely unaffected, potentially preserving nuclear basket function.

To further investigate the mechanism underlying the loss of degradative activity in SPOP-F102C compared with SPOP-F133L, we performed affinity purification-mass spectrometry (AP-MS) using SPOP as the bait (**Figure S2** and **Table S3**). The results showed that both SPOP-WT and SPOP-F133L interacted with NUP153, whereas SPOP-F102C did not (**Figure 2C** and **Figure S2**), suggesting that the impaired degradation of NUP153 by SPOP-F102C may result from the loss of physical interaction. This difference likely reflects the distinct biochemical consequences of the two amino-acid substitutions, namely that substitution of phenylalanine with cysteine at position 102 introduces a smaller and more polar side chain that disrupts the local hydrophobic packing critical for substrate recognition, whereas substitution with leucine at position 133 may largely preserves hydrophobicity and side-chain volume, causing only minimal perturbation of the NUP153-binding interface. These findings highlight an interesting avenue for future structural studies on the molecular basis of SPOP-substrate interactions. Our AP-MS did not detect TPR, possibly due to weak or transient binding between SPOP and TPR. Future work employing approaches optimized to capture such transient interactions could provide further insight. In addition, the AP-MS identified NUP50 as another interactor of SPOP-WT and SPOP-F133L, but not SPOP-F102C (**Figure 2C** and **Figure S2**). This interaction did not lead to substrate degradation in the whole-proteome analysis (**Figure 2B**), consistent with prior reports that SPOP binds but does not degrade NUP50^[16]^. Moreover, these interactions identified by AP-MS were validated by co-immunoprecipitation using HA-tagged SPOP constructs (**Figure 2D**).

### p53 is down-regulated in SPOP-F133L-expressing cells through a CRL-dependent post-translational mechanism

Despite being generally considered loss-of-function mutations, our data suggest that SPOP-F133L may retain degradative activity toward certain substrates. We also considered the possibility that these mutants might acquire gain-of-function properties by recognizing and degrading neo-substrates. To investigate this, we sought to identify proteins that are specifically down-regulated in cells expressing SPOP mutants but not in those expressing SPOP-WT.

Because SPOP regulates a number of transcriptional factors and co-regulators^[17]^, resulting in broad proteomic alterations, changes observed in whole-proteome analyses could arise from either transcriptional or post-translational effects. Therefore, we focused on functionally important proteins that showed differential expressions and aimed to validate whether these changes occur through post-translational regulation. Among these candidates, p53 caught our attention as a well-characterized tumor suppressor and transcription factor that regulates cell-cycle arrest, DNA repair, apoptosis, and senescence in response to cellular stress^[18]^. Notably, SPOP-WT and SPOP-F102C did not alter p53 protein levels, whereas SPOP-F133L expression led to modest down-regulation of p53 (**Figure 3A,B**). This observation was further confirmed by Western blot analysis (**Figure 3C**). Given that SPOP functions as part of a CRL E3 complex requiring Cullin neddylation for activity, we employed MLN4924, a selective inhibitor that blocks CRL activation by preventing Cullin neddylation^[19]^, to assess whether the down-regulation of p53 depends on CRL activity. Treatment with MLN4924 rescued the p53 down-regulation induced by SPOP-F133L (**Figure 3D**), indicating that this effect occurs via a post-translational mechanism mediated by a CRL E3 ligase.

**Figure 3.**
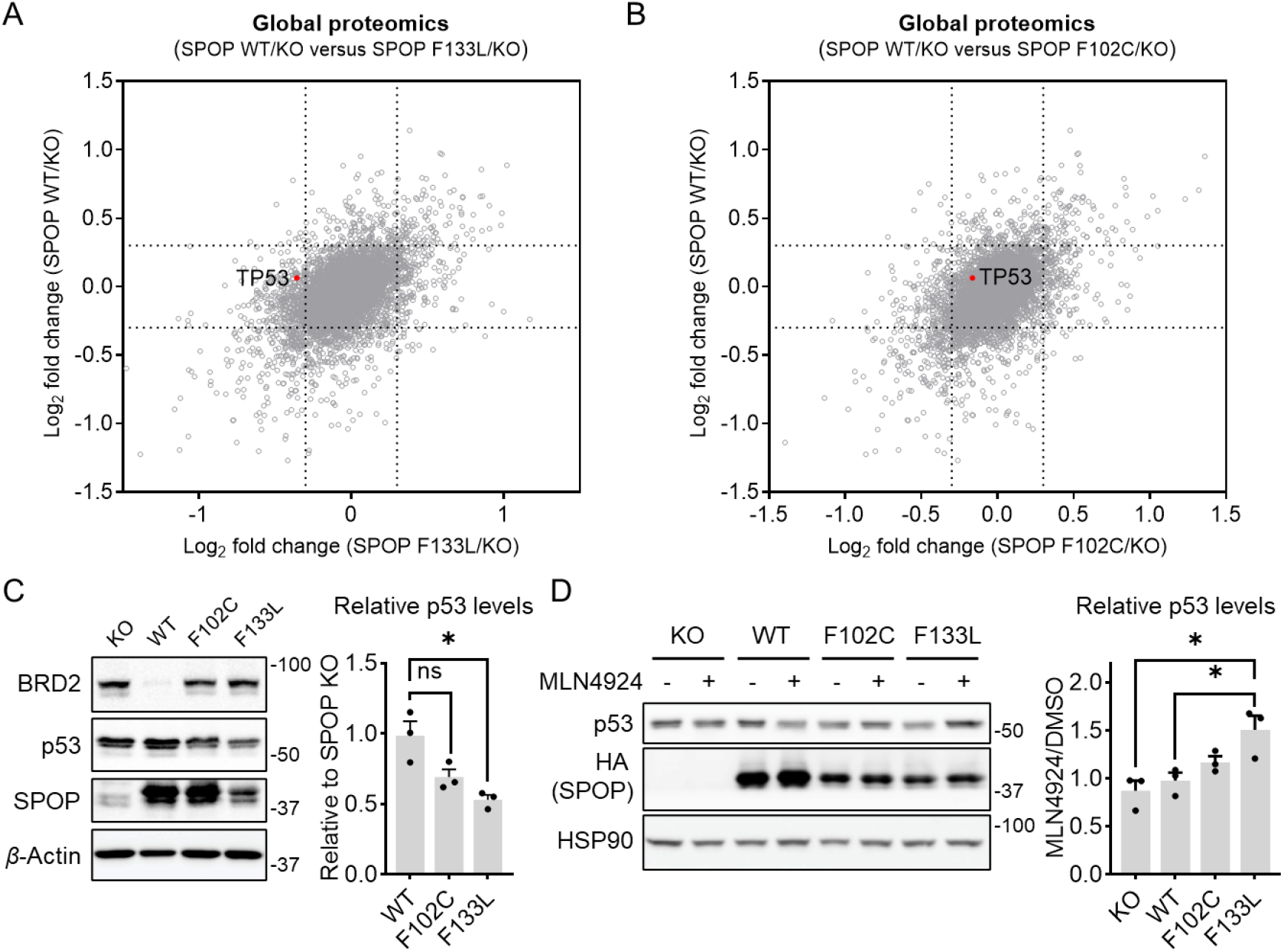
p53 is down-regulated in SPOP-F133L-expressing cells through a CRL-dependent post-translational mechanism. **A**. Global proteomic analysis comparing two calculated ratios, SPOP-WT/KO and SPOP-F133L/KO, to identify proteins differentially affected by the SPOP-F133L mutant. **B**. Global proteomic analysis comparing two calculated ratios, SPOP-WT/KO and SPOP-F102C/KO, to identify proteins differentially affected by the SPOP-F102C mutant. **C**. Western blot analysis of p53 expression in *SPOP* KO HEK293T cells and in KO cells stably expressing SPOP-WT, F102C, or F133L. BRD2 serves as a positive control. The result is representative of three experiments. The bar graph represents quantification of p53 protein levels normalized to *β*-Actin. Data are presented as mean ± SEM (n = 3 biological independent samples). **D**. Western blot analysis of p53 expression in *SPOP* KO HEK293T cells and in KO cells stably expressing SPOP-WT, F102C, or F133, with or without treatment with 2 μM MLN4924 for 4 hours. The result is representative of three experiments. The bar graph shows the quantification of p53 protein levels (normalized to *β*-Actin) in MLN4924-treated versus DMSO-treated cells. Data are presented as mean ± SEM (n = 3 biological independent samples).

p53 is a master regulator of various pathways that suppress tumorigenesis^[18]^. To determine whether modest down-regulation of p53 by SPOP-F133L leads to measurable phenotypic changes, we assessed several p53-dependent cellular processes, including cell proliferation, sensitivity to chemotherapeutic agents, and DNA damage-induced cell-cycle arrest, in cells expressing SPOP-WT, SPOP-F102C, or SPOP-F133L. We first monitored cell proliferation over 96 hours and observed no significant differences in growth rate among the four cell models (**Figure S3A**). We next tested cellular responses to doxorubicin, a chemotherapeutic agent that induces DNA double-strand breaks and oxidative stress, thereby activating the p53 pathway and triggering cell-cycle arrest and apoptosis^[20]^. In principle, cells with reduced p53 levels may show increased resistance to doxorubicin-induced apoptosis. However, cytotoxicity assays revealed no significant shift in IC_50_ values between SPOP-WT and SPOP-F133L-expressing cells (**Figure S3B**). Similarly, cell-cycle arrest assays following doxorubicin treatment showed no significant differences among these cells **(Figure S3C**,**D**). We speculate that two explanations may explain the lack of detectable phenotypic changes following p53 down-regulation. First, the extent of down-regulation was relatively mild, with p53 protein levels reduced by 22% in SPOP-F133L-expressing cells compared with *SPOP* KO cells (*p* value = 0.005), which may be insufficient to elicit strong alterations in p53-related pathways. Second, HEK293T cells may not represent an optimal system for evaluating tumorigenic phenotypes. Nonetheless, our findings highlight a potentially new regulatory mechanism in SPOP-F133L-expressing cells, thereby providing a basis for future investigation of this pathway in additional cancer-relevant models.

### SPOP-F102C and SPOP-F133L are TPD-competent E3 ligases

TPD has emerged as a powerful approach to manipulate protein homeostasis. By using small molecules or biologics to link target proteins to endogenous degradation systems such as the proteasome or lysosome, TPD enables their selective and efficient removal^[21]^. SPOP is one of the several TPD-competent E3 ligases and has been demonstrated to support biologics-mediated protein degradation^[22]^. We recently introduced the concept of exploiting the somatic mutant FBXW7-R465C for targeted protein degradation, highlighting the potential to develop cancer-selective degraders^[23]^. Building on this idea, we asked whether SPOP mutants might similarly serve as TPD-competent ligases. As a proof-of-concept, we generated HaloTag-fused SPOP-WT and mutant constructs (**Figure 4A**) and synthesized JQ1-HL, a bifunctional compound that links a HaloTag-reactive chloroalkane ligand^[24]^ to the BRD4 inhibitor JQ1^[25]^ (**Figure 4B**). This system allows us to assess BRD4 degradation as well as the resulting cytotoxicity, given that BRD4 is an essential gene whose loss can induce cell death^[26]^.

**Figure 4.**
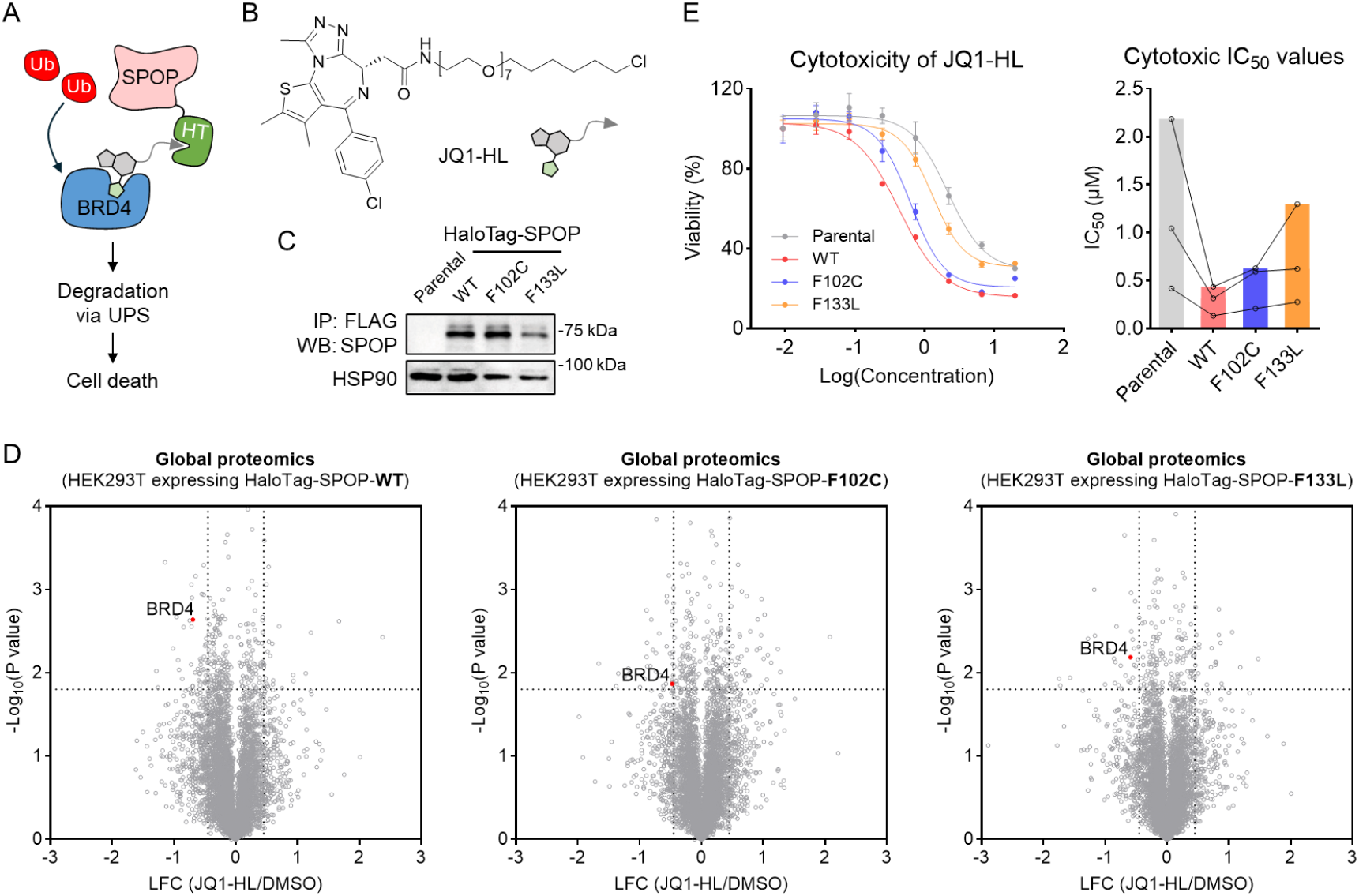
SPOP-F102C and SPOP-F133L support targeted protein degradation. **A**. Schematic illustration of SPOP-mediated BRD4 degradation, in which HaloTag (HT)-fused SPOP and a bifunctional degrader cooperatively recruit BRD4 for proteasomal degradation. **B**. Structure of JQ1-HL. **C**. Western blot analysis of SPOP expression in parental HEK293T cells and in HEK293T cells stably expressing HaloTag-SPOP-WT, F102C, or F133L.The result is representative of two experiments (n = 2 biologically independent samples). **D**. Volcano plot showing global proteomic changes in HEK293T cells stably expressing HaloTag-SPOP-WT, F102C, or F133L, with or without JQ1-HL treatment (1 µM, 24 hours). Protein expression in HaloTag-SPOP-expressing cells treated with JQ1-HL was normalized to the BRD4 ratio in HEK293T parental cells treated with JQ1-HL versus DMSO. *P* values were calculated by two-sided t-test and adjusted using Benjamini-Hochberg correction for multiple comparisons (n = 2 biological independent samples). **E**. Dose-response curves and IC_50_ values of JQ1-HL-induced cytotoxicity in parental HEK293T cells and HEK293T cells stably expressing HaloTag-SPOP-WT, F102C, or F133L. Data are presented as mean ± SEM (n = 3 biological independent samples).

We established HEK293T cell lines stably expressing HaloTag-SPOP-WT, -F102C, or - F133L (**Figure 4C**) and performed whole-proteome analysis following treatment with JQ1-HL. In all three models, JQ1-HL treatment induced significant BRD4 degradation (**Figure 4D** and **Table S4**). Furthermore, this degradation translated into enhanced cytotoxicity, as reflected by decreased IC_50_ values for JQ1-HL in cells expressing HaloTag-SPOP (**Figure 4E**). Together, these findings suggest that SPOP-F102C and SPOP-F133L are capable of supporting targeted protein degradation. Future studies exploring both heterobifunctional and molecular glue degraders may yield new ligands that hijack these SPOP mutants to drive selective target degradation in tumors harboring these mutations.

## Conclusion

In this study, we combined systematic proteomic and functional analyses to reveal that SPOP-F133L, but not SPOP-F102C, retains degradative activity toward the nuclear basket proteins NUP153 and TPR, and can partially reduce p53 levels through a CRL-dependent post-translational mechanism. In addition, both mutants were able to support targeted protein degradation in an engineered cellular system, suggesting that they may retain functional competence in mediating ligand-induced degradation despite altered substrate recognition. Together, these findings expand our understanding of SPOP mutation-associated functional diversity and suggest that certain cancer-associated variants might be repurposed as TPD-competent E3 ligases for therapeutic applications.

## Acknowledgements

We gratefully acknowledge the support of the NIH R35GM154945 (X.Z.), the Falk Medical Research Trust Catalyst Award (X.Z.), and the Carcinogenesis Training Program at the Northwestern University Feinberg School of Medicine (A.C.).

## Data Availability Statement

The mass spectrometry proteomics data have been deposited to the ProteomeXchange Consortium via the PRIDE^[27]^ partner repository with the dataset identifier PXD071054.

## References

[1] G. A. Collins, A. L. Goldberg, Cell 2017, 169, 792–806.

[2] N. Zheng, N. Shabek, Annu Rev Biochem 2017, 86, 129–157.

[3] M. D. Petroski, R. J. Deshaies, Nat Rev Mol Cell Biol 2005, 6, 9–20.

[4] M. Zhuang, M. F. Calabrese, J. Liu, M. B. Waddell, A. Nourse, M. Hammel, D. J. Miller, H. Walden, D. M. Duda, S. N. Seyedin, T. Hoggard, J. W. Harper, K. P. White, B. A. Schulman, Mol Cell 2009, 36, 39–50.

[5] M. R. Marzahn, S. Marada, J. Lee, A. Nourse, S. Kenrick, H. Zhao, G. Ben-Nissan, R. M. Kolaitis, J. L. Peters, S. Pounds, W. J. Errington, G. G. Prive, J. P. Taylor, M. Sharon, P. Schuck, S. K. Ogden, T. Mittag, EMBO J 2016, 35, 1254–1275.

[6] X. Dai, W. Gan, X. Li, S. Wang, W. Zhang, L. Huang, S. Liu, Q. Zhong, J. Guo, J. Zhang, T. Chen, K. Shimizu, F. Beca, M. Blattner, D. Vasudevan, D. L. Buckley, J. Qi, L. Buser, P. Liu, H. Inuzuka, A. H. Beck, L. Wang, P. J. Wild, L. A. Garraway, M. A. Rubin, C. E. Barbieri, K. K. Wong, S. K. Muthuswamy, J. Huang, Y. Chen, J. E. Bradner, W. Wei, Nat Med 2017, 23, 1063–1071.

[7] J. An, C. Wang, Y. Deng, L. Yu, H. Huang, Cell Rep 2014, 6, 657–669.

[8] A. C. Groner, L. Cato, J. de Tribolet-Hardy, T. Bernasocchi, H. Janouskova, D. Melchers, R. Houtman, A. C. B. Cato, P. Tschopp, L. Gu, A. Corsinotti, Q. Zhong, C. Fankhauser, C. Fritz, C. Poyet, U. Wagner, T. Guo, R. Aebersold, L. A. Garraway, P. J. Wild, J. P. Theurillat, M. Brown, Cancer Cell 2016, 29, 846–858.

[9] aC. Kandoth, M. D. McLellan, F. Vandin, K. Ye, B. Niu, C. Lu, M. Xie, Q. Zhang, J. F. McMichael, M. A. Wyczalkowski, M. D. M. Leiserson, C. A. Miller, J. S. Welch, M. J. Walter, M. C. Wendl, T. J. Ley, R. K. Wilson, B. J. Raphael, L. Ding, Nature 2013, 502, 333–339; bC. E. Barbieri, S. C. Baca, M. S. Lawrence, F. Demichelis, M. Blattner, J. P. Theurillat, T. A. White, P. Stojanov, E. Van Allen, N. Stransky, E. Nickerson, S. S. Chae, G. Boysen, D. Auclair, R. C. Onofrio, K. Park, N. Kitabayashi, T. Y. MacDonald, K. Sheikh, T. Vuong, C. Guiducci, K. Cibulskis, A. Sivachenko, S. L. Carter, G. Saksena, D. Voet, W. M. Hussain, A. H. Ramos, W. Winckler, M. C. Redman, K. Ardlie, A. K. Tewari, J. M. Mosquera, N. Rupp, P. J. Wild, H. Moch, C. Morrissey, P. S. Nelson, P. W. Kantoff, S. B. Gabriel, T. R. Golub, M. Meyerson, E. S. Lander, G. Getz, M. A. Rubin, L. A. Garraway, Nat Genet 2012, 44, 685–689.

[10] M. Blattner, D. J. Lee, C. O’Reilly, K. Park, T. Y. MacDonald, F. Khani, K. R. Turner, Y. L. Chiu, P. J. Wild, I. Dolgalev, A. Heguy, A. Sboner, S. Ramazangolu, H. Hieronymus, C. Sawyers, A. K. Tewari, H. Moch, G. S. Yoon, Y. C. Known, O. Andren, K. Fall, F. Demichelis, J. M. Mosquera, B. D. Robinson, C. E. Barbieri, M. A. Rubin, Neoplasia 2014, 16, 14–20.

[11] M. Nakazawa, M. Fang, H. M. C. T. L. Lotan, P. Isaacsson Velho, E. S. Antonarakis, Prostate 2022, 82, 260–268.

[12] Y. Deng, W. Ding, K. Ma, M. Zhan, L. Sun, Z. Zhou, L. Lu, Cell Death Dis 2024, 15, 172.

[13] aM. Blattner, D. Liu, B. D. Robinson, D. Huang, A. Poliakov, D. Gao, S. Nataraj, L. D. Deonarine, M. A. Augello, V. Sailer, L. Ponnala, M. Ittmann, A. M. Chinnaiyan, A. Sboner, Y. Chen, M. A. Rubin, C. E. Barbieri, Cancer Cell 2017, 31, 436–451; bW. Gan, X. Dai, A. Lunardi, Z. Li, H. Inuzuka, P. Liu, S. Varmeh, J. Zhang, L. Cheng, Y. Sun, J. M. Asara, A. H. Beck, J. Huang, P. P. Pandolfi, W. Wei, Mol Cell 2015, 59, 917–930; cJ. P. Theurillat, N. D. Udeshi, W. J. Errington, T. Svinkina, S. C. Baca, M. Pop, P. J. Wild, M. Blattner, A. C. Groner, M. A. Rubin, H. Moch, G. G. Prive, S. A. Carr, L. A. Garraway, Science 2014, 346, 85–89.

[14] aC. Geng, B. He, L. Xu, C. E. Barbieri, V. K. Eedunuri, S. A. Chew, M. Zimmermann, R. Bond, J. Shou, C. Li, M. Blattner, D. M. Lonard, F. Demichelis, C. Coarfa, M. A. Rubin, P. Zhou, B. W. O’Malley, N. Mitsiades, Proc Natl Acad Sci U S A 2013, 110, 6997–7002; bH. Janouskova, G. El Tekle, E. Bellini, N. D. Udeshi, A. Rinaldi, A. Ulbricht, T. Bernasocchi, G. Civenni, M. Losa, T. Svinkina, C. M. Bielski, G. V. Kryukov, L. Cascione, S. Napoli, R. I. Enchev, D. G. Mutch, M. E. Carney, A. Berchuck, B. J. N. Winterhoff, R. R. Broaddus, P. Schraml, H. Moch, F. Bertoni, C. V. Catapano, M. Peter, S. A. Carr, L. A. Garraway, P. J. Wild, J. P. Theurillat, Nat Med 2017, 23, 1046–1054.

[15] M. Beck, E. Hurt, Nat Rev Mol Cell Biol 2017, 18, 73–89.

[16] J. Y. Ong, M. Abdusamad, I. Ramirez, A. Gholkar, X. Zhang, T. V. Gimeno, J. Z. Torres, Mol Biol Cell 2025, 36, ar24.

[17] A. Clark, M. Burleson, Am J Cancer Res 2020, 10, 704–726.

[18] E. R. Kastenhuber, S. W. Lowe, Cell 2017, 170, 1062–1078.

[19] T. A. Soucy, P. G. Smith, M. A. Milhollen, A. J. Berger, J. M. Gavin, S. Adhikari, J. E. Brownell, K. E. Burke, D. P. Cardin, S. Critchley, C. A. Cullis, A. Doucette, J. J. Garnsey, J. L. Gaulin, R. E. Gershman, A. R. Lublinsky, A. McDonald, H. Mizutani, U. Narayanan, E. J. Olhava, S. Peluso, M. Rezaei, M. D. Sintchak, T. Talreja, M. P. Thomas, T. Traore, S. Vyskocil, G. S. Weatherhead, J. Yu, J. Zhang, L. R. Dick, C. F. Claiborne, M. Rolfe, J. B. Bolen, S. P. Langston, Nature 2009, 458, 732–736.

[20] C. Carvalho, R. X. Santos, S. Cardoso, S. Correia, P. J. Oliveira, M. S. Santos, P. I. Moreira, Curr Med Chem 2009, 16, 3267–3285.

[21] aM. Schapira, M. F. Calabrese, A. N. Bullock, C. M. Crews, Nat Rev Drug Discov 2019, 18, 949–963; bJ. A. Wells, K. Kumru, Nat Rev Drug Discov 2024, 23, 126–140.

[22] aY. J. Shin, S. K. Park, Y. J. Jung, Y. N. Kim, K. S. Kim, O. K. Park, S. H. Kwon, S. H. Jeon, A. Trinh le, S. E. Fraser, Y. Kee, B. J. Hwang, Sci Rep 2015, 5, 14269; bS. C. Chang, P. Gopal, S. Lim, X. Wei, A. Chandramohan, R. Mangadu, J. Smith, S. Ng, M. Gindy, U. Phan, B. Henry, A. W. Partridge, Cell Chem Biol 2022, 29, 1601–1615 e1607; cM. Hoffman, D. Krum, K. D. Wittrup, J Biol Chem 2024, 300, 107616; dS. Lim, R. Khoo, Y. C. Juang, P. Gopal, H. Zhang, C. Yeo, K. M. Peh, J. Teo, S. Ng, B. Henry, A. W. Partridge, ACS Cent Sci 2021, 7, 274–291; eS. Lim, R. Khoo, K. M. Peh, J. Teo, S. C. Chang, S. Ng, G. L. Beilhartz, R. A. Melnyk, C. W. Johannes, C. J. Brown, D. P. Lane, B. Henry, A. W. Partridge, Proc Natl Acad Sci U S A 2020, 117, 5791–5800.

[23] A. A. Basu, C. Zhang, M. Rouhimoghadam, A. Vasudevan, J. M. Reitsma, X. Zhang, J Am Chem Soc 2025, 147, 6108–6115.

[24] G. V. Los, L. P. Encell, M. G. McDougall, D. D. Hartzell, N. Karassina, C. Zimprich, M. G. Wood, R. Learish, R. F. Ohana, M. Urh, D. Simpson, J. Mendez, K. Zimmerman, P. Otto, G. Vidugiris, J. Zhu, A. Darzins, D. H. Klaubert, R. F. Bulleit, K. V. Wood, ACS Chem Biol 2008, 3, 373–382.

[25] P. Filippakopoulos, J. Qi, S. Picaud, Y. Shen, W. B. Smith, O. Fedorov, E. M. Morse, T. Keates, T. T. Hickman, I. Felletar, M. Philpott, S. Munro, M. R. McKeown, Y. Wang, A. L. Christie, N. West, M. J. Cameron, B. Schwartz, T. D. Heightman, N. La Thangue, C. A. French, O. Wiest, A. L. Kung, S. Knapp, J. E. Bradner, Nature 2010, 468, 1067–1073.

[26] R. Arafeh, T. Shibue, J. M. Dempster, W. C. Hahn, F. Vazquez, Nat Rev Cancer 2025, 25, 59–73.

[27] Y. Perez-Riverol, C. Bandla, D. J. Kundu, S. Kamatchinathan, J. Bai, S. Hewapathirana, N. S. John, A. Prakash, M. Walzer, S. Wang, J. A. Vizcaino, Nucleic Acids Res 2025, 53, D543–D553.

