## Supporting Information for "Systematic characterization of cancer-associated SPOP mutants reveals novel and reprogrammable degradative activities"

### Supporting Tables

Supplementary Table 1: Global proteomics comparing protein expression in HEK293T parental and *SPOP* KO cells.

Supplementary Table 2: Global proteomics comparing protein expression in *SPOP* KO HEK293T cells and KO cells stably expressing SPOP-WT, F102C, or F133L.

Supplementary Table 3: AP-MS analysis of the SPOP interactome in HEK293T cells expressing HA-tagged SPOP-WT, F102C, or F133L.

Supplementary Table 4: Global proteomics comparing protein expression in HEK293T cells stably expressing HaloTag-SPOP-WT, F102C, or F133L, with or without JQ1-HL treatment.

**Supplementary Figures**

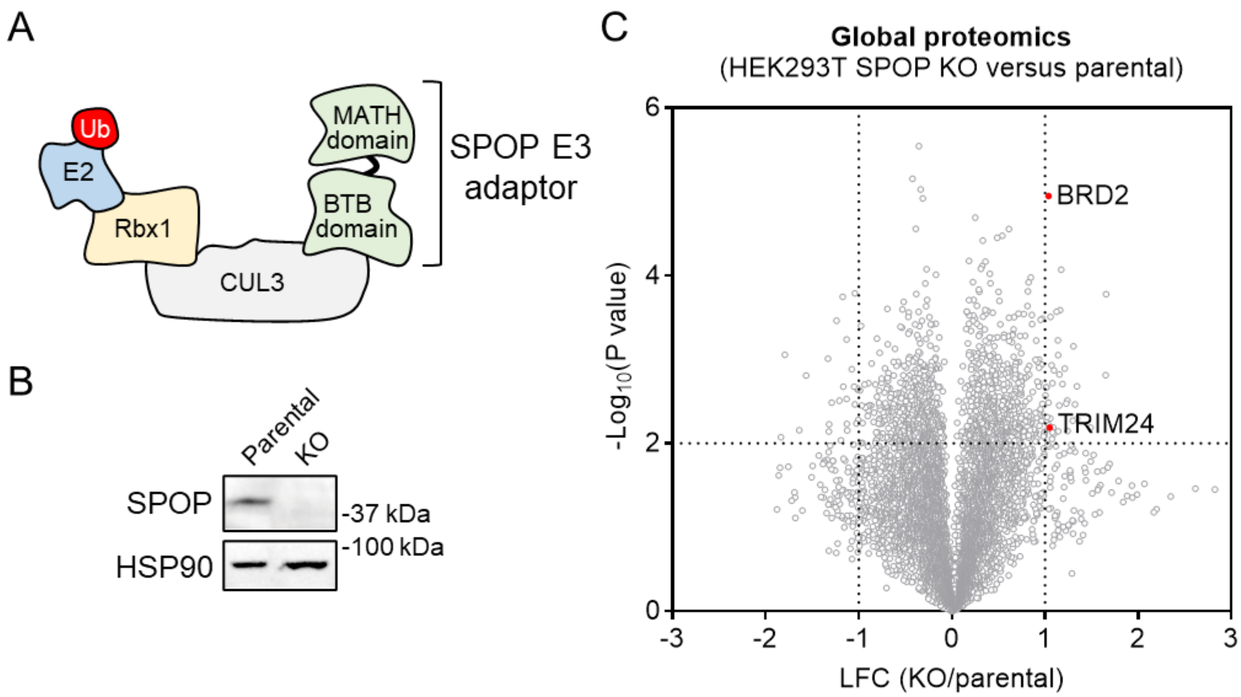


**Figure S1. Generation and characterization of *SPOP* KO HEK293T cells. A**. Schematic representation of the SPOP-CUL3-Rbx1 E3 ligase complex. **B**. Western blot analysis of SPOP expression in parental and *SPOP* KO HEK293T cells. The result is representative of two experiments (n = 2 biologically independent samples). **C**. Volcano plot showing global proteomic changes between parental and *SPOP* KO HEK293T cells (n = 4 biological independent samples for parental cells, n = 3 biological independent samples for *SPOP* KO cells). *P* values were calculated by two-sided *t*-test and adjusted using Benjamini-Hochberg correction for multiple comparisons. LFC, $log_2$ fold change.

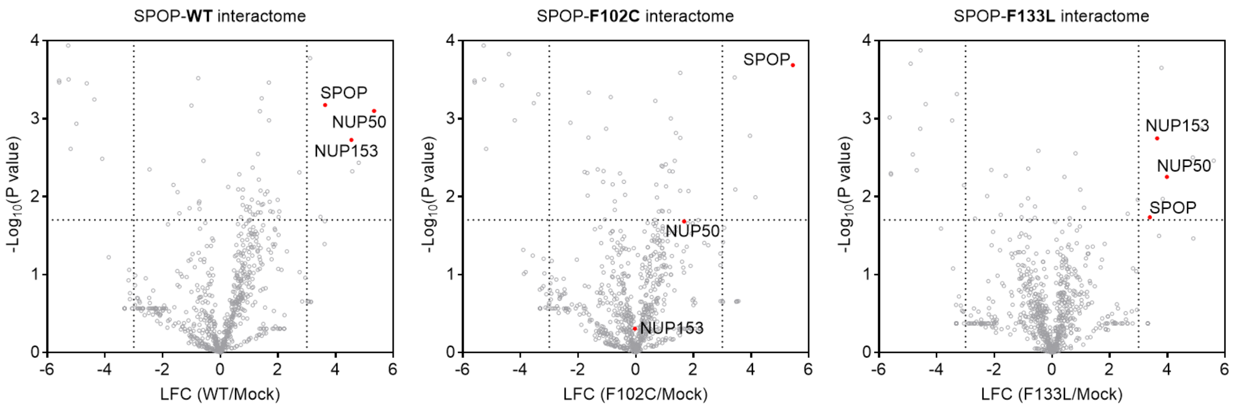


**Figure S2**. Volcano plot showing the SPOP interactome compared with mock-transfected cells (n = 2 biologically independent samples for mock and SPOP-F133L, n = 3 biologically independent samples for SPOP-WT and F102C). *P* values were determined using a two-sided t-test and adjusted for multiple comparisons using the Benjamini-Hochberg method.

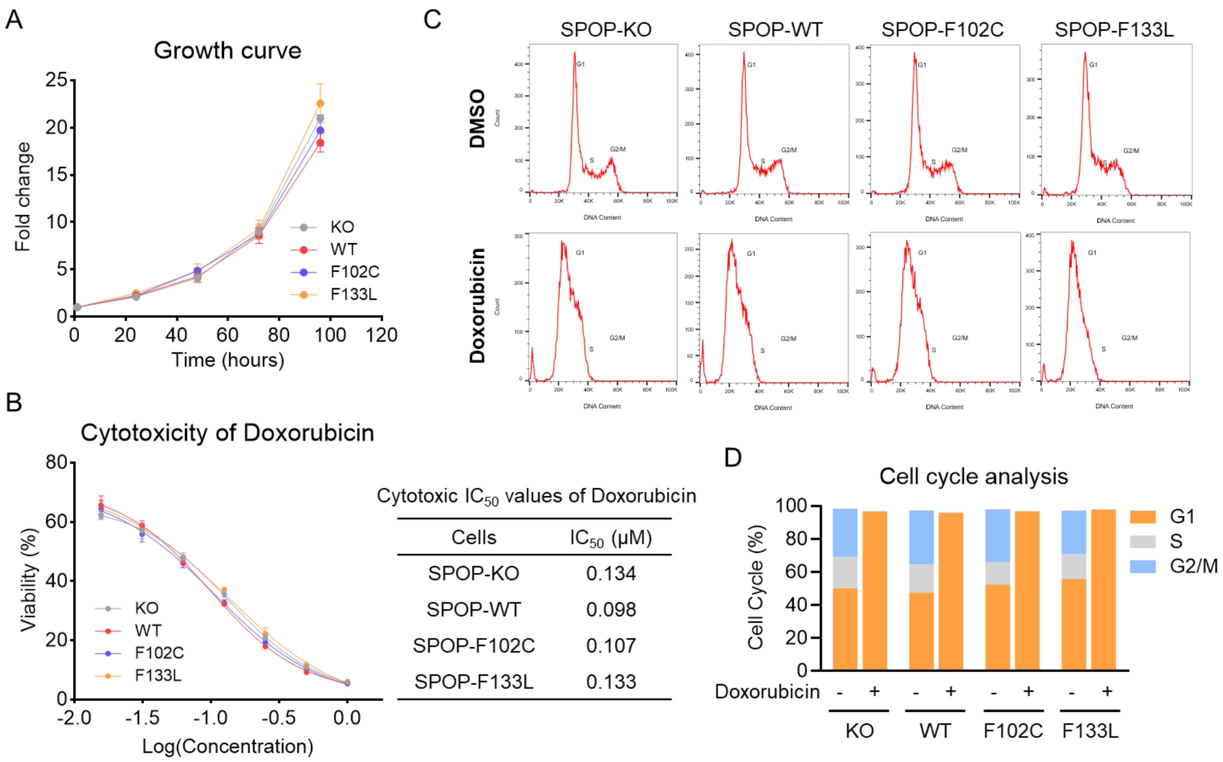


Cytotoxic $IC_{50}$ values of Doxorubicin

| Cells | $IC_{50}$ (μM) |
|---|---|
| SPOP-KO | 0.134 |
| SPOP-WT | 0.098 |
| SPOP-F102C | 0.107 |
| SPOP-F133L | 0.133 |

**Figure S3. Partial down-regulation of p53 by SPOP-F133L does not produce detectable phenotypic changes. A**. Cell proliferation curves of *SPOP* KO HEK293T cells and KO cells stably expressing SPOP-WT, F102C, or F133L, measured over 96 hours by cell viability assays. Data are presented as mean ± SEM (n = 3). **B**. Dose-response curves showing cell viability following treatment with Doxorubicin. $IC_{50}$ values were calculated using nonlinear regression analysis. Data are presented as mean ± SEM (n = 3). **C**. Representative flow cytometry profiles of cell-cycle distribution following treatment with Doxorubicin (1 μM, 24 hours). **D**. Quantification of cell-cycle phases ($G_1$, S, and $G_2$/M) from (C) showing no significant differences among the four cell models.

### Materials and Methods

#### Reagents

Anti-FLAG affinity gel (clone M2, cat. #A2220), anti-HA HRP antibody (clone 3F10, cat. #12013819001), and cOmplete protease inhibitor cocktail (cat. #11873580001) were purchased from Sigma-Aldrich. Anti-HA Sepharose beads (clone C29F4, cat. #3956), anti-BRD2 antibody (clone D89B4, cat. #5848), anti-p53 antibody (clone 1C12, cat. #2524), anti-NUP153 antibody (clone E3I6Z, cat. #98559), anti-HSP90 HRP antibody (clone C45G5, cat. #79641), and anti-β-Actin HRP antibody (clone 13E5, cat. #5125) were purchased from Cell Signaling Technology. Anti-SPOP antibody (clone 2B9F2, cat. #68216-1-Ig) and anti-NUP50 antibody (clone 1D5A7, cat. #67001-1-Ig) were purchased from Proteintech. Cas9 endonuclease was purchased from Integrated DNA Technologies (cat. #1081061). FuGene 6 (cat. #E2692) transfection reagent and sequencing grade trypsin/Lys-C mix (cat. #V5071) were purchased from Promega. MLN4924 (cat. #15217) was purchased from Cayman Chemical. Enzyme-linked chemiluminescence (ECL) (cat. #32106), ECL plus (cat. #32132), BCA protein assay kit (cat. #23227), and Tandem Mass Tag (TMT) isobaric label reagent (cat. #90110) were purchased from Thermo Scientific.

#### Molecular Cloning

Human SPOP with an N-terminal HA tag was purchased as a gene block from Integrated DNA Technologies and cloned into the pCDH-CMV-MCS-EF1-Blast vector using the NheI and NotI restriction sites. SPOP amino acid substitutions were generated using the Q5 site-directed mutagenesis kit (New England Biolabs) with primers designed via NEBaseChanger.

#### Cell Culture and Generation of Stable Expression Cells

HEK293T cells were obtained from ATCC. HEK293T cells were maintained in Dulbecco’s Modified Eagle Medium (DMEM, Corning) supplemented with 10% fetal bovine serum (vol/vol), 2 mM L-glutamine (Gibco), and Penicillin/Streptomycin at 37°C in 5% $CO_2$. Stable expression of SPOP variants and Halo-tagged SPOP variants was achieved by lentiviral transduction. Lentivirus was produced by co-transfecting the plasmid containing gene of interest with psPAX2 and pMD2.G into HEK293T cells using FuGene 6. Viral supernatant was collected 48 and 72 hours post-transfection and filtered through a 0.45 µm Millex-HV sterile syringe filter unit (MilliporeSigma). For lentiviral transduction, 100,000 cells were seeded in a 6-well plate and incubated overnight. The following day, 1 mL of lentiviral supernatant was added together with 8 µg/mL Polybrene. After 6 hours of incubation, the virus-containing medium was replaced with complete DMEM. After 48 hours, cells were selected with 10 µg/mL Blasticidin or 2 µg/mL Puromycin.

#### Generation of CRISPR-Cas9 *SPOP* Knockout Cells

HEK293T cells with *SPOP* knockout were generated by electroporating Cas9-sgRNA ribonucleoprotein complexes using the 4D-Nucleofector system (Lonza Bioscience). Two sgRNAs targeting the SPOP gene were combined for electroporation (sgRNA #1: CCACTCGACATTTCTGCCGG; sgRNA #2: GTTATTGATGGTCCACATGT).

#### Cell Viability Assay

Cells were plated in 96-well clear-bottom white plates (Corning) at a density of 3,000 cells per well in 100 µL of complete DMEM, followed by addition of 1 µL of compound. Plates were incubated for 72 hours. For cell-growth curves, cells were plated in 96-well plates at 3,000 cells per well in 100 µL of complete DMEM and incubated under standard conditions. After incubation, plates were equilibrated to room temperature for 15 minutes, and 50 µL of CellTiter-Glo reagent (Promega) was added to each well. Plates were incubated for 15 minutes at room temperature, and luminescence was measured using a CLARIOstar Plus microplate reader (BMG Labtech).

#### Flow Cytometry

Cells were plated in 10 cm dishes and allowed to adhere overnight. The next day, 1 μM doxorubicin or an equivalent volume of DMSO was added, and cells were incubated for 24 hours at 37°C. After treatment, cells were collected and 1 million cells were transferred to a fresh 15 mL conical tube. Hoechst stain was added at two drops per milliliter of medium, and samples were incubated for 1 hour at 37°C. Cells were then pelleted, resuspended in 1 mL FACS buffer, and passed through a filter tube to remove clumps. Samples were analyzed on a BD Fortessa flow cytometer, and cell-cycle distribution was determined using FlowJo software.

**Co-Immunoprecipitation**

Samples were lysed in 500 μL NP-40 lysis buffer supplemented with cOmplete EDTA-free Protease Inhibitor Cocktail (1 tablet per 10 mL; Sigma-Aldrich) and 200 U/mL Pierce Universal Nuclease for Cell Lysis (Fisher Scientific). Protein concentrations were measured using the DC assay, and lysates were diluted to 1 mg/mL in 500 μL lysis buffer. A 25 μL aliquot was reserved as the input sample. Lysates were incubated with either anti-FLAG or anti-HA beads at 4°C with rotation for 2 hours. Beads were washed four times with immunoprecipitation wash buffer (0.2% NP-40, 25 mM Tris-HCl pH 7.4, 150 mM NaCl) and once with PBS. Samples were then analyzed by western blot or subjected to affinity-purification mass spectrometry.

**Affinity Purification Mass Spectrometry**

Following immunoprecipitation, beads were resuspended in 40 μL of 8 M urea and incubated at 65°C for 10 minutes to denature proteins. Proteins were eluted using a BioSpin column (Bio-Rad). Samples were reduced with 5 μL of 200 mM dithiothreitol and incubated at 65°C for 15 minutes. Alkylation was performed by adding 5 μL of 400 mM iodoacetamide and shaking the samples at 37°C in the dark for 30 minutes. To dilute the urea concentration, 300 μL PBS was added to each sample, followed by 2 μg trypsin/Lys-C mix. Digestion was carried out at 37°C with shaking for 18 hours. TMT labeling was performed by adding 5 μL of TMT reagent to each sample and incubating at room temperature for 1 hour. Reactions were quenched with 6 µL of 5% hydroxylamine for 15 minutes at room temperature, after which 4 µL formic acid was added. Samples were dried in a speed vacuum and desalted using a Sep-Pak C18 Cartridge (Waters).

Peptides were dissolved in buffer A (4% acetonitrile with 0.1% formic acid) and injected onto a Vanquish Neo UHPLC system coupled to an Orbitrap Eclipse Tribrid Mass Spectrometer. Peptide separation was performed on an EASY-Spray PepMap Neo C18 column (2 µm, 75 µm × 150 mm; Thermo Scientific, ES75150PN) at a flow rate of 0.25 µL/min. Chromatographic elution was achieved using solvent A (0.1% formic acid in water) and solvent B (80% acetonitrile with 0.1% formic acid) with the following gradient: 5% B for the first 15 minutes, a linear increase from 5% to 35% B between 15-165 minutes, ramping to 100% B from 165-170 minutes, and held at 100% B for an additional 10 minutes. The spray voltage on the nano-ESI source was set to 2 kV. MS1 spectra were recorded in the Orbitrap at a resolution of 60,000 and m/z range of 400-1600, with the RF lens set to 30%, a normalized AGC target of 100%, and a maximum injection time of 118 ms. Precursor ions were isolated with a 0.7 m/z window and fragmented using HCD in the ion trap (normalized AGC target 100%; collision energy 30%; maximum injection time 35 ms). For SPS-MS3 acquisition, up to 10 MS2 fragment ions were selected for additional HCD fragmentation, and MS3 scans were acquired in the Orbitrap (collision energy 55%; normalized AGC target 500%; maximum injection time 118 ms; resolution 60,000; m/z 100-500). Data acquisition was performed using Xcalibur v4.7.102.25.

Raw data files were processed with Proteome Discoverer 2.5 (Thermo Scientific). The Sequest HT search engine was used to search against the UniProt human reference proteome (UP000005640_9606_Human.fasta) using fully tryptic specificity, a precursor mass tolerance of 10 ppm, and a fragment mass tolerance of 0.6 Da. Carbamidomethylation of cysteine residues and TMT tags on peptide N-termini and lysines were specified as fixed modifications, whereas methionine oxidation and protein N-terminal acetylation were included as variable modifications. Peptide-spectrum matches were rescored with Percolator and filtered to a 1% FDR at both the peptide and protein levels. Reporter ion intensities were obtained from SPS-MS3 scans using a 20-

ppm integration window, and quantification was performed using unique and razor peptides with total peptide-signal normalization across all channels.

### Global Proteomics

Cells were lysed in 80 µL PBS using sonication (5 pulses at 40% intensity, 3 rounds). Protein concentrations were determined using the DC assay, and lysates were diluted to 1 mg/mL. For each sample, 50 µL (equivalent to 50 µg total protein) was denatured in 12 M urea. For reduction, 5 µL of a 200 mM dithiothreitol stock solution was added and samples were incubated at 65°C for 15 minutes. Alkylation was performed by adding 5 µL of a 400 mM iodoacetamide stock solution and shaking the samples at 37°C in the dark for 30 minutes. Urea concentration was reduced by adding 300 µL PBS to each sample, followed by addition of 2 µg trypsin/Lys-C mix. Digestion proceeded at 37°C for 18 hours. For TMT labeling, 35 µL of each digested peptide sample was used. To each sample, 9 µL of acetonitrile was added, followed by 3 µL of TMT reagent. Samples were incubated at room temperature for 1 hour. Labeling reactions were quenched by adding 6 µL of 5% hydroxylamine, followed by 2.5 µL formic acid. Labeled samples were pooled and dried in a speed vacuum.

The combined sample was resuspended in 300 µL of 0.1% (vol/vol) formic acid in water. High-pH reversed-phase fractionation was performed using the High pH Reversed-Phase Peptide Fractionation Kit (Fisher Scientific) to generate 30 fractions. These 30 fractions were combined into 10 final fractions by pooling fractions 1, 11, and 21; 2, 12, and 22; and so on through 10, 20, and 30. Fractions were dried by speed vacuum, resuspended in 20 µL of 0.1% formic acid, sonicated for 5 minutes, centrifuged, and analyzed by LC-MS/MS following the same data acquisition and analysis procedures described in the Affinity Purification Mass Spectrometry section.

### Statistical Analysis

Quantitative data are presented as scatter plots with the mean and standard error of the mean (SEM) shown as error bars. Comparisons between two groups were performed using an unpaired two-tailed Student's t-test. Statistical significance is indicated as follows: *$P$ < 0.05, **$P$ < 0.01, ***$P$ < 0.001, and ****$P$ < 0.0001.

**Synthetic Procedures**

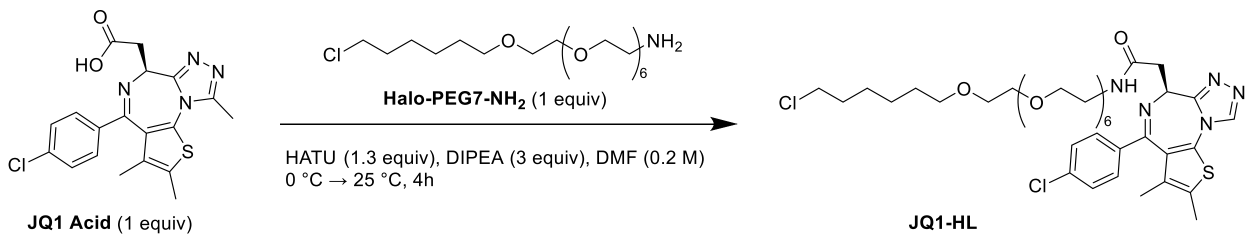


**(*R*)-*N*-(27-chloro-3,6,9,12,15,18,21-heptaoxaheptacosyl)-2-(4-(4-chlorophenyl)-2,3,9-trimethyl-6*H*-thieno[3,2-f][1,2,4]triazolo[4,3-a][1,4]diazepin-6-yl)acetamide (JQ1-HL)**

The synthesis of Halo-PEG7-$NH_2$ was performed according to a previously reported procedure[1]. JQ1 Acid (30 mg, 0.067 mmol, 1 eq), Halo-PEG7-$NH_2$ (27.1 mg, 0.067 mmol, 1 eq), and HATU (33.4 mg, 0.087 mmol, 1.3 eq) was dissolved in DMF (1 mL) on ice. The DIPEA (35.4 µL, 0.201 mmol, 3 eq) was added and stirred at room temperature for 2 hours. Upon completion, the reaction was concentrated and purified by flash chromatography to yield JQ1-HL as yellow oil (17.8 mg, 0.021 mmol, 32%).

**$^1$H NMR (500 MHz, $CDCl_3$)**: δ 7.40 (d, $J$ = 8.5 Hz, 2H), 7.31 (d, $J$ = 8.5 Hz, 2H), 6.90 (t, $J$ = 5.5 Hz, 1H), 4.64 (t, $J$ = 7.0 Hz, 1H), 3.69-3.48 (m, 30H),3.39-3.33 (m, 1H), 2.65 (s, 3H), 2.38 (s, 3H), 1.75 (m, 2H), 1.57 (m, 2H), 1.48 – 1.30 (m, 4H), 1.24 (s, 6H).

**$^{13}$C NMR (126 MHz, $CDCl_3$)**: δ 170.42, 163.63, 155.53, 149.67, 136.56, 136.54, 132.05, 130.75, 130.57, 130.31, 129.71(2C), 128.53(2C), 71.07-70.22(m, 9C), 69.93, 69.69, 54.22, 44.91, 39.28, 38.97, 32.40, 29.54, 29.30, 26.55, 25.27, 14.25, 12.93, 11.69.

**HRMS (ESI⁺)** m/z calcd for $C_{39}H_{57}Cl_2N_5O_8S$ $[M+H]^+$: 848.3203; found 848.3238.

**NMR Spectra**

[1]H NMR of JQ1-HL

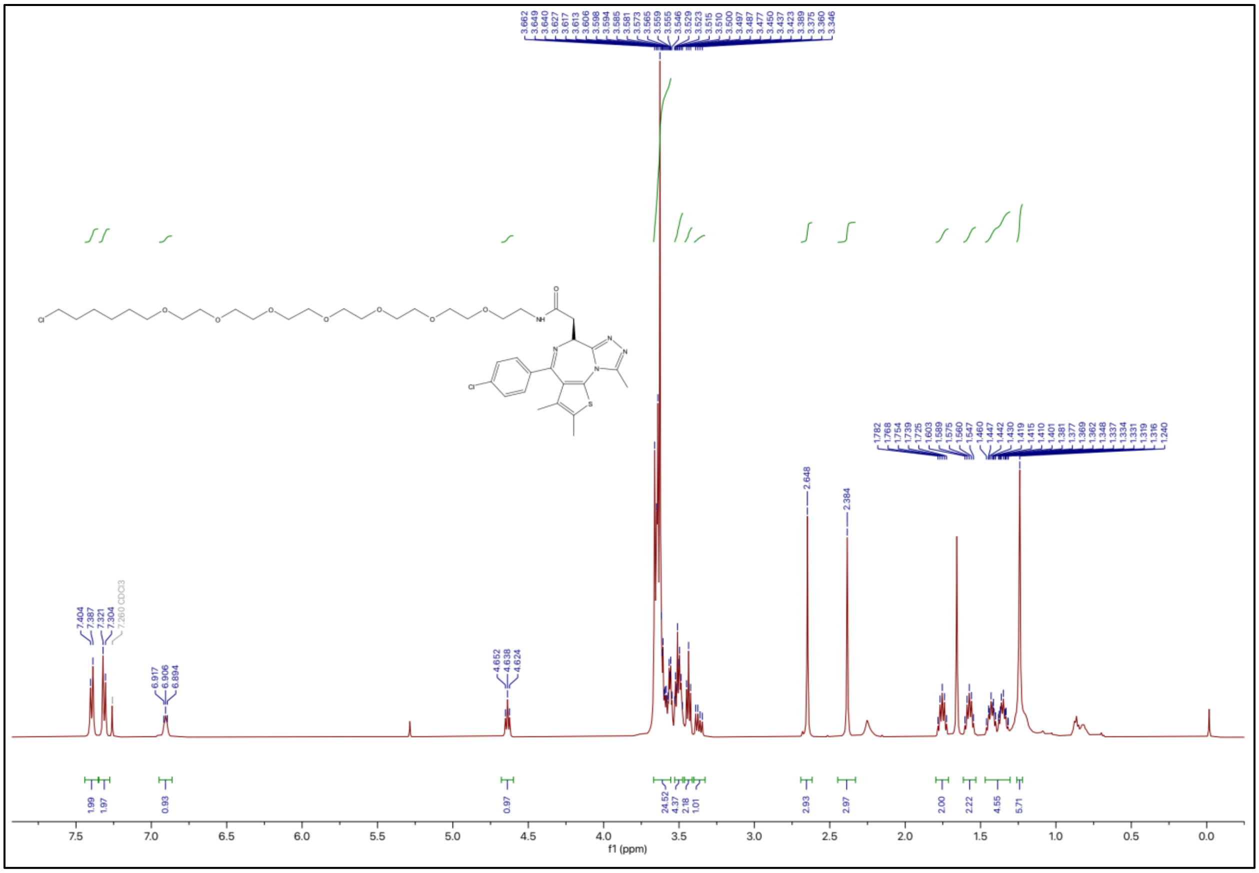

$^{13}C$ NMR of JQ1-HL

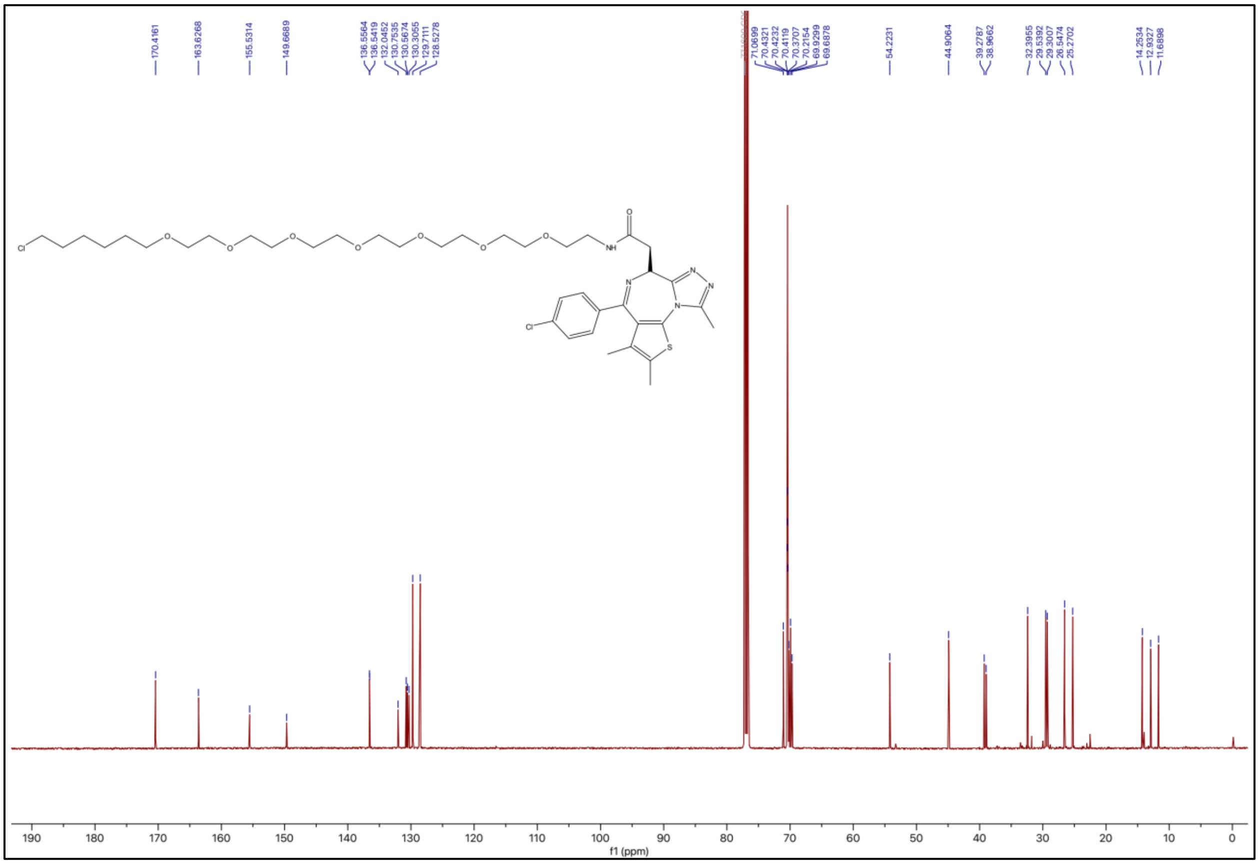


### References


[1] Z. Hu, P. H. Chen, W. Li, T. Douglas, J. Hines, Y. Liu, C. M. Crews, *J Am Chem Soc* **2023**.
